# Land Use Cover changes in the western escarpment of Rift Valley in the Gamo Zone, Southern Ethiopia

**DOI:** 10.1101/2021.09.08.459379

**Authors:** Temesgen Dingamo, Serekebirhan Takele, Sebsebe Demissew, Zerihun Woldu

## Abstract

LULC changes are caused by natural and human alterations of the landscape that could largely affect forest biodiversity and the environment. The aim of the study was to analyzed LULC change dynamics in the western escarpment of the rift valley of the Gamo Zone, Southern Ethiopia. Digital satellite images downloaded from USGS were analyzed using ERDAS Imagine (14) and Arc GIS 10.2 software and supervised image classification was used to generate LULC classification, accuracy assessment and Normalized Difference Vegetation Index (NDVI). Drivers of LULC change were identified and analyzed. Four land classes were identified such as forest, farmland, settlement and water-wetland. Settlement and farmlands have increased by 7.83% and 5.88%, respectively. On the other hand, both forest and water bodies and wetland decreased by aerial coverage of 11.03% and 2.68%, respectively. The overall accuracy of the study area was 92.86%, 94.22% and 94.3% with a kappa value of 0.902, 0.92 and 0.922, respectively. NDVI values ranged between -0.42 to 0.73. Agricultural expansion (31.4%), expansion of settlement (25.7%) and Fuelwood collection and Charcoal production (22.9%) were the main driving forces that jeopardize forest biodiversity of the study area. Integrated land use and policy to protect biodiversity loss, forest degradation and climate changes are deemed necessary.

## 1. INTRODUCTION

Land use land cover change (LULCC) is a major issue of concern with regards to change in a global environment [1]; changes are so pervasive such that, when aggregated globally, they significantly affect key aspects of Earth System functioning[2,3]. This directly impacts biodiversity throughout the world [4]; contribute to local and regional climate change [5] as well as to global climate warming [6]; are the primary sources of soil degradation [7]; and, by altering ecosystem services, affect the ability of biological systems supporting human needs [8]. Such changes also determine, in part, the vulnerability of places and people to climatic, economic, or socio-political perturbations [9].

The land is the major natural resource in which economic, social, infrastructure and other human activities are undertaken [10]. Thus, changes in land use that has occurred at all times in the past, currently on-going, and is likely to continue in the future [11, 12]. These changes have beneficial or detrimental impacts, the latter being the principal causes of global concern as they impact human well-being and safety [13; 3]. LULC changes are widespread, accelerating, and the trade-offs offset human livelihood [14]. The rapid growth and expansion of urban centers, population pressure, scarcity of land, changing technologies are among the many drivers of LULC in the world today [15].

[16] Stated that land cover change occurs through conversion and intensification by human intervention, altering the balance of an ecosystem, generating a response expressed as system changes. For centuries, humans have been altering the earth’s surface to produce food through agricultural activities [17]. In the past few decades, the conversion of grasslands, woodlands, and forests into croplands and pastures has risen dramatically, especially in developing countries where a large proportion of the human population depends on natural resources for their livelihoods [17, 18, and 19]. The increasing demand for land and related resources often results in changes in land use/cover [16] and it has local, national, regional and global causes and implications [20].

In Africa, forests cover about (21.4%) of the land area which corresponds to 674 million hectares and in Eastern Africa alone approximately 13% of the land area is under forests and woodlands [21]. [22] noted that close to 40% of Ethiopia might have been covered by high forests and that about 16% of the land area was covered by high forests in the early 1950s (EFAP 1994). In the early 1980s the high forest cover of Ethiopia declined to 3.6% and further declined to 2.7 % in 1989 [23]. The recent estimate of the land cover of Ethiopia that could qualify as ‘forests’ which includes high forests, woodlands, plantations, and bamboo forests adds up to 15% [24].

Land cover change occurs naturally in a progressive manner but, could sometimes be rapid and abrupt due to anthropogenic activities [25]. Vegetation cover change is a process in which the level of diversity and the density of individual species that makes up the natural vegetation structure are altered as a result of natural and human-induced pressure [26; 27]. Vegetation change mapping and monitoring are useful when changes in the vegetation attributes of interest result in detectable changes in image radiance, emittance, or microwave backscatter values [28]. Many research results in Ethiopia indicate some of the critical threats to forests that need to be seriously addressed. One of these is land use/ cover changes [29, 30 and 31]. There is a dearth of LULC change detection studies in the study area and hence, the present study aims to evaluate and analyze LULC change detection at the southwest escarpment of the rift valley of Gamo Zone, Southern Ethiopia.

## 2. MATERIALS and METHODS

### 2.1 Description of the Study Area

The study was carried out in the western escarpment of the rift valley of the Gamo Zone, Southern Ethiopia. Gamo Zone is bordered by Dirashe Special Woreda in the South, Gofa Zone in the NW, Dawro and Wolayita Zones in the north, Lake Abaya and Chamo in the NE, South Omo in the South and Amaro Special Woreda in the SE (Figure 1). Araba Minch town is the administrative center of Gamo Zone.

**Figure 1:**
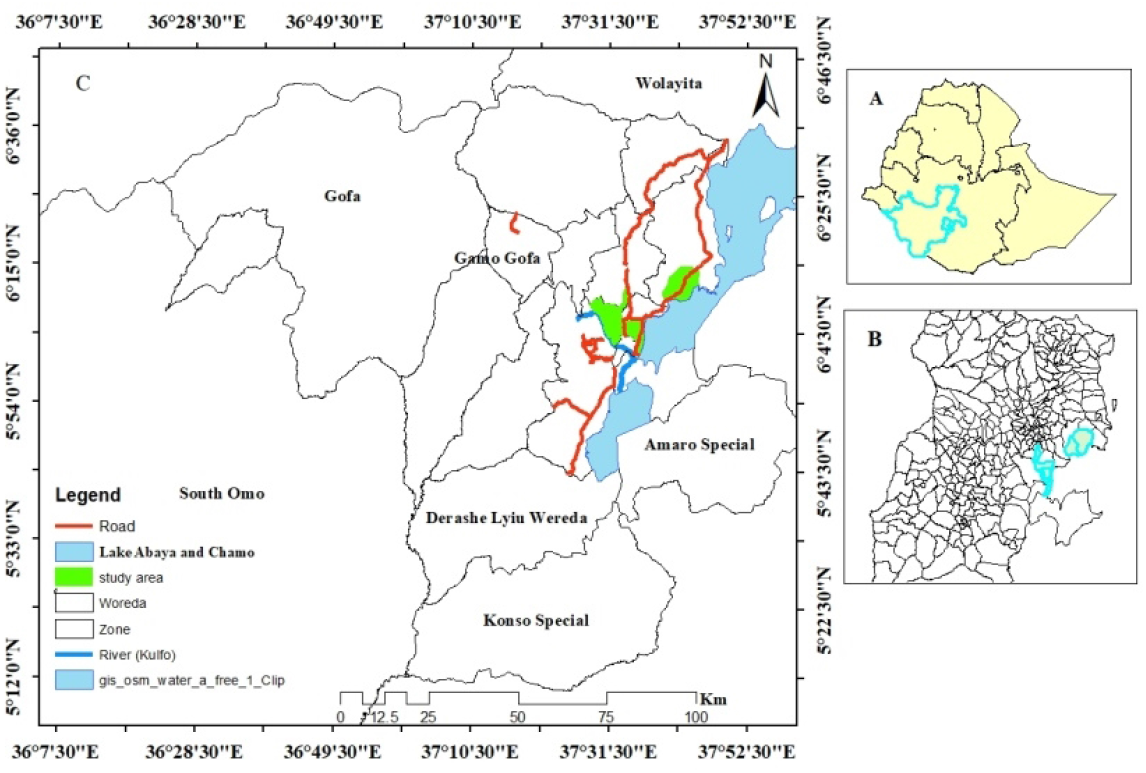
Location map for the study area (A = Ethio-Region, B = Gamo Zone, C =Study area (surrounded Zone, Lake Abaya and Chamo, Rivers and Roads all weathered)) (Source: Arc GIS 10.2 and CSA)

The study area consists of plains and hillsides of the Gamo mountain ridge between 6°05’N to 6°12’N and 37°33’E to 37°39’E. The elevation of the area ranging from 1168 m to 2535 m a.s.l and the slope of the forest ranges between 0 to 32 degrees (Figure 2). The total population in the study area is estimated to be 195,858 in the 2019 projection population (CSA, 2019) (Table 1). Drainage in the study area is seasonal and many streams from the mountain chains merge to form the Kulfo and Hara rivers which eventually join the western escarpment of the Central Rift Valley to Lakes (Chamo and Abaya).

**Figure 2:**
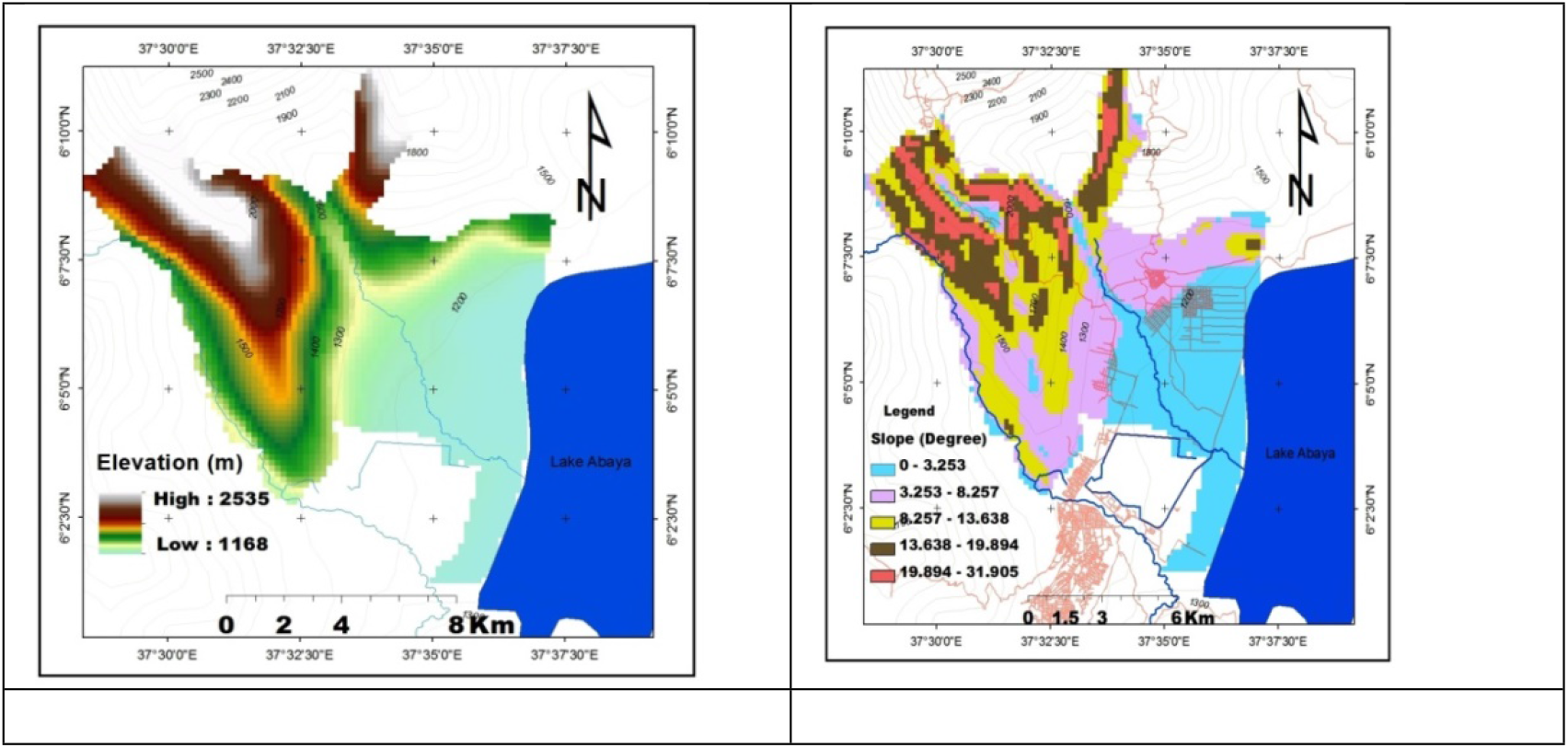
Elevation and Slope Map

**Table 1:** Total Population of the study area from 1999-2019 (CSA, 2019)

| Year | Arba Minch Zuria |  |  |  |
| --- | --- | --- | --- | --- |
|  | M | F | Total |  |
| 1999 | 58,062 | 55,468 | 113,530 |  |
| 2009 | 82,751 | 82,929 | 165,680 |  |
| 2019 | 97,905 | 97,953 | 195,858 | projection population |

#### Population

The total population of the study area was increased in the three successive periods (1999, 2009 and 2019) (Table 1) (CSA, 2019).

#### Geology and soil

The geology of the Rift-valley escarpment is mainly quaternary volcanic alluvial deposits and lacustrine clay. Forest and the state farm are composed of three main types: Fluvisols, Gleysols and Vertisols. Fluvisols consist of soil materials developed in alluvial deposits and flood plains [32]. The Rift valley floor near Lake Abaya and Chamo is filled with alluvial sediments. The bedrock in the region consists of basalt, trachyte, rhyolite, and ignimbrite and the western edges of Lake Abaya are covered by approximately 1 to 2-km wide plain of lacustrine and swamp deposits [33]. The topsoil textural classes of major soils in its spatial distribution are mainly dominated by clay loam, light clay, loam sand and sandy clay loam based on USDA classification.

#### Vegetation Cover

According to [34], the study area is characterized by complex vegetation types such as *Combretum-Terminalia* woodland vegetation, *Acacia-Commiphora* woodland vegetation and Dry evergreen Montana forest. The most common tree species in the study area are *Terminalia brownii, Combretum molle, Ziziphus mucronata, Pappea capensis, Cadaba farinosa, Vachellia and Senegalia Acacia species, Balanites aegyptiaca, Commiphora abyssinica, Rhus natalensis, Olea europaea, Psydrax schimperiana, Acokanthera schimperi, etc*.

#### Climate

The study area has a bimodal rainfall type. Maximum and minimum mean annual rainfall during 1999-2019 was 1141.1 mm and 491.8 mm, respectively (Figure 3). The maximum and minimum mean annual temperature was 33.6^0^C and 15^0^C, respectively (Figure 3) [35].

**Figure 3:**
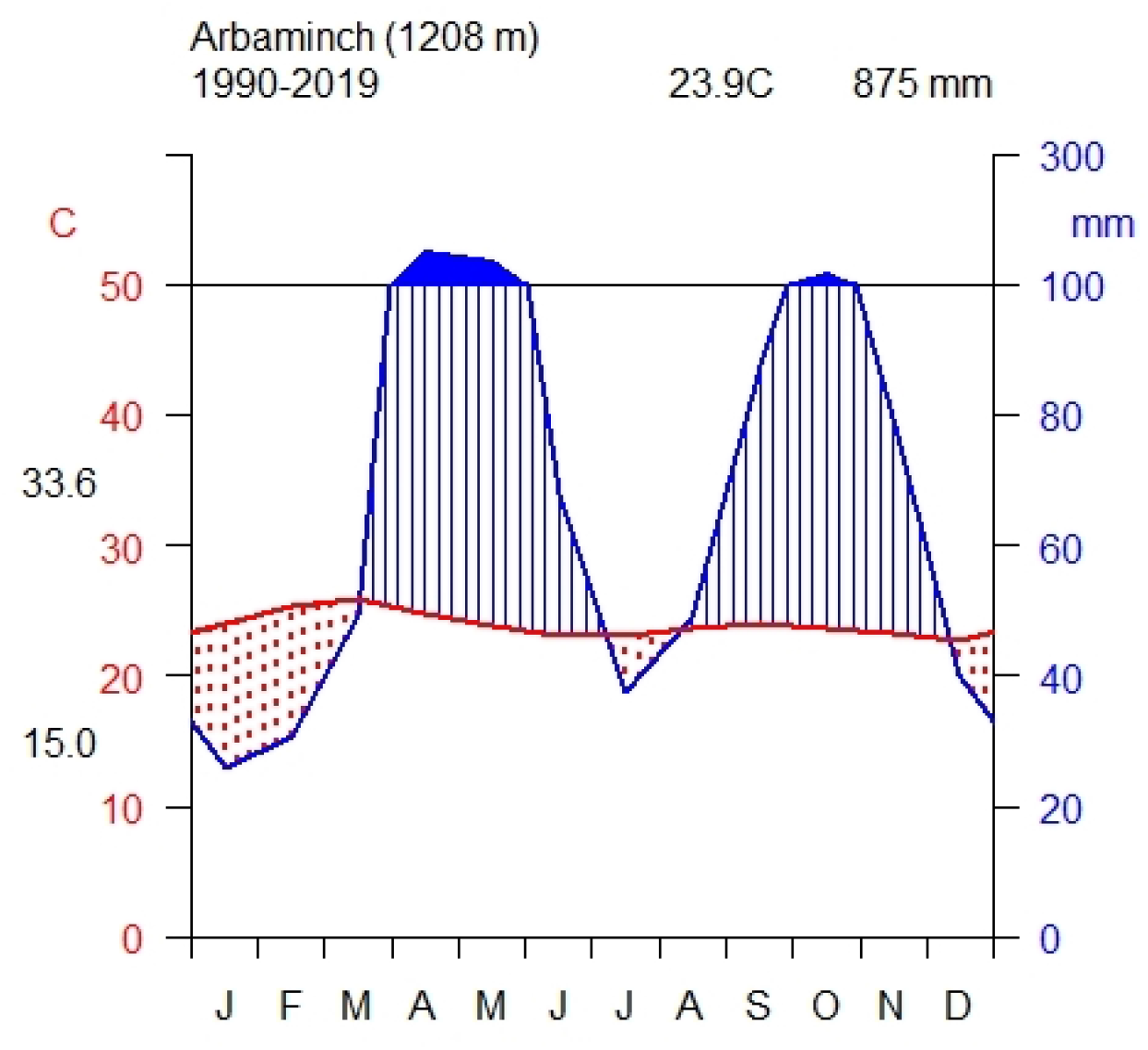
Annual Max. and Min. temp.in **°C** and rainfall in mm (1990-2019)

### 2.2 Data types and sources

Primary and secondary data were used: Ground control points (GCP) for ground truth were collected as primary data using handheld GPS. Secondary data include Landsat Thematic Mapper (TM) for the year 1999, ETM+ for the year 2009 and Landsat 8 Operational Land Imager (OLI) images for the year 2019 acquired from United States Geological Survey online imagery portals (http://glovis.usgs.gov). Other Geo-spatial data include Shapefiles and topographic maps collected from the Central Statistical Agency (CSA) and Ethiopian Mapping Agency (EMA) for extraction and delineation of area of interest (Table 2).

**Table 2:** Remote sensing data of the study

| Acquisition data | Sensors | Path and Row | Spatial Resolution | Number of bands | Format | Source |
| --- | --- | --- | --- | --- | --- | --- |
| 01/05/1999 | TM | 169, 56 | 30m | 7 | TIFF | USGS |
| 01/05/2009 | ETM<sup>+</sup> | 169,56 | 30m | 8 | TIFF | USGS |
| 01/05/2019 | OLI | 169,56 | 30m | 11 | TIFF | USGS |

### 2.3 Land-use change assessment (1999–2019)

Digital satellite images were processed classified and analyzed using ERDAS Imagine (14). Computations of the area and changes in land use categories were made using Arc GIS 10.2 software analytical tools. Pre-processing of satellite images was done to create a more faithful representation of the original scene. An intensive pre-processing such as geo-referencing, layer-stacking, resolution merge, and sub sets were carried out to Ortho-rectify the satellite images into UTM coordinates (WGS, 1984) and to remove disturbances such as haze, noise, steep slope effect, and radiometric variation between acquisition dates. A stacked satellite image of the study area was extracted by clipping the Area of Interest (AOI) layer of the Gamo shapefile in ERDAS 14 software.

The satellite image was classified using the **s**upervised image classification technique and employed pixel-based supervised image classifications with the maximum likelihood classification algorithm [36] to produce LULC maps of the study area. Appropriate band combinations were obtained and the signatures were used for the supervised classification. Land cover change detection for the study area was monitored at three intervals: 1999_2009, 2009_2019 and 1999_2019. Supervised classification into four land classes were categories and distinguished into farmlands, forest lands, settlement, water bodies and wetlands (Table 3).

**Table 3:** Characteristics of land cover classes

| Class name | Description |
| --- | --- |
| Farmlands | Areas used for crop cultivation (Maze, teff, Banana, Mango, etc.). |
| Dense forest<br>scattered forest<br>and woodland | This habitat is dominated by trees characterized by a multi-storeyed nature with the crown cover of almost 10-50% |
| Settlement | Different settlements (villages) associated with building |
| Water_wetland | areas where water cover and may support both aquatic and wetland species |

### 2.4 Accuracy analysis

Since image classification without accuracy assessment is incomplete [37], accuracy assessment for the images was carried out. The accuracy of the classification was assessed using producers, users and overall methods of accuracy assessment. The overall accuracy, as well as Kappa statics, was calculated based on the GCP collected from the identified land-use types. Kappa statics was calculated by the following equation:-

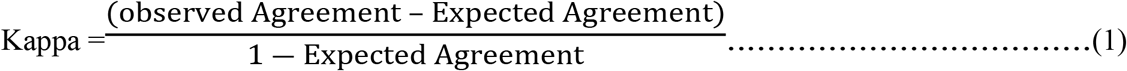

### 2.5 Land use land covers change detection

The LULC maps of three years showing period’s with a range of ten years in between (1999, 2009 and 2019) were generated from the satellite imageries using supervised maximum likelihood classification. To analyze the land cover structural changes in the study area the table showing the area in hectares and percentage changes between the periods 1999_2009, 2009_2019 and 1999-2019 were measured for each LULC type. Change detection was calculated by:-

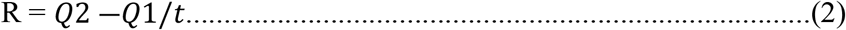

Where, R = Rate of Change, Q_2_ = Recent year forest cover in ha

Q_1_ = Initial Year forest cover in ha and

t = Interval year between Initial year and Recent year

### 2.6 Vegetation index

Normalized Difference Vegetation Index (NDVI) is one of the indicators commonly used to detect the vegetation cover of the earth’s surface i.e. spectral change detection method. NDVI values were calculated on composite image and used band 3 (Red) and 4 (Near Infrared) for Landsat 7, and band 4 (Red) come with band 5 (Near Infrared) for Landsat 8. NDVI approaching calculation of greenness degree of image correlates with vegetation crown density. NDVI correlates with chlorophyll content and its value is between -1 to 1. NDVI is calculated as:

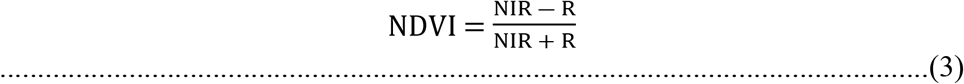

Where: NDVI = Normalized Difference Vegetation Index, NIR=Near Infra-Red Band R= Red Band

### 2.7 Drivers of LULC changes

LULC changes are influenced by a number of driving factors. In the study area, human activity is often mentioned as the major driver of LULC Changes. For a better understanding of LULC changes data were collected including field observation, focused group discussion (FDG) and key informant interview (KII). KII and FGD were selected based on the recommendation of local community leaders and agriculture extension workers. The participants included elders (male and female), agriculture extension workers and youth jobless. The informants were asked for their consent to participate in the discussion were then given clear information about LULC changes in the study area. Data were analyzed using IBM SPSS version 20.

## 3. RESULTS AND DISCUSSION

### 3.1 Land use land covers classification

The Four land classes identified in the study include forest, farmland, settlement and water bodies and water-wetlands. The land use land cover categories in Figure 4 show that forest land class has progressively decreased while farmlands and settlement increased from 1999_2019.

**Figure 4:**
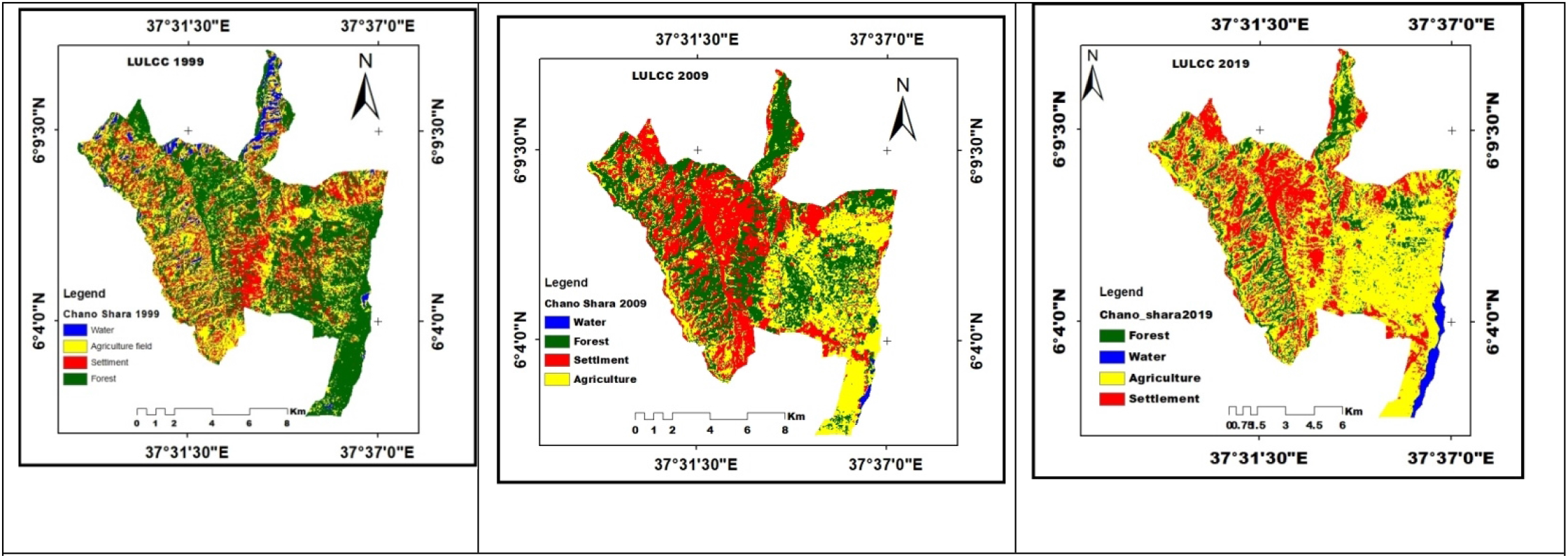
Land use land cover change from 1999_2019

Similar results were reported by [38; 39; and 40] showing that farmlands in the Rift Valley of Ethiopia have expended as a result of population pressure.[41] has shown that more than 4/5 of the total terrestrial productive land in the Ethiopian Central Rift Valley was lost to agriculture. Conversions to other land use types have been observed and the image classification shows a clear conversion of land covers into farmland and settlement (Table 4).

**Table 4:** Land use land covers change (1999_2019)

| Area | Area (ha)<br>( 1999_2009) | % | Area (ha)<br>( 2009_2019) | % | Area (ha)<br>(1999-2019) | % |
| --- | --- | --- | --- | --- | --- | --- |
| Forest - Agriculture | 1416.156 | 11.4 | 2671.36 | 21.51 | 1878.42 | 15.13 |
| Forest - Settlement | 500.144 | 4.03 | 284.27 | 2.29 | 376.529 | 3.03 |
| Agriculture - Forest | 155.922 | 1.26 | 105.00 | 0.85 | 50.315 | 0.41 |
| Agriculture-Settlement | 142.651 | 1.15 | 408.7 | 3.29 | 376.529 | 3.03 |
| Agriculture - Water | 235.9683 | 1.9 | 166.1 | 1.34 | 232.268 | 1.87 |
| Water - Agriculture | 384.342 | 3.09 | 401.85 | 3.24 | 277.00 | 2.23 |

### 3.2 Land use land covers change

Results revealed that the extent of land cover changes from forest to farmland in the last three decades was rapid. The decline of water bodies and wetlands was not as dramatic as the loss of forests (Table 4). The conversion of farmlands to settlements was equally high. Similar results were reported by [42 and 43] in the Finchaa Catchment, North-western Ethiopia and Abijata Shalla National Park, respectively. This is due to small-scale irrigation by pumping water from the lakes and rivers for income generation through the production of fruit and vegetables. [44] also showed that urban settlements and farmland expansion gained the most in the area compared to other LULC types, while forest areas exhibited a decreasing trend. Demand for food and grazing land for the growing population appears to be the driving factors.

### 3.3 Land use land covers change detection

LULC change detection was showing that the areal coverage of settlement and farmlands increased. On the other hand, both forest and water_wetland were decreased by an aerial coverage (Table 5). This was due to the conversion of forest and water_wetland, to settlement and farmlands increased and also Lake Abaya might be fluctuated increased and or decreased its volume, but mostly at the expense of forest lands (Table 5). [44] shown that urban settlements and farmland expansion gained the most in the area compared to other LULC types, while forest areas exhibited a decreasing trend (Figure 5). Demand for food and grazing land for the growing population seems the probable driving force, among others.

**Table 5:** Land use land covers change detection from 1999 to 2019

| Land class | Rate change (r=Q2-Q1/t ) |  |  |  |  |  |
| --- | --- | --- | --- | --- | --- | --- |
|  | 1999_2009 |  | 2009_2019 |  | 1999_2019 |  |
|  | ha | % | ha | % | ha | % |
| Settlement | 574.17 | 4.6 | 398.45 | 3.2 | 972.63 | 7.83 |
| Agriculture | 628.62 | 5.06 | 101.1 | 0.81 | 729.72 | 5.88 |
| Forest | -199.95 | -1.61 | -1169.79 | -9.42 | -1369.74 | -11.03 |
| Water | -1002.83 | -8.08 | 670.23 | 5.4 | -332.6 | -2.68 |

**Figure 5:**
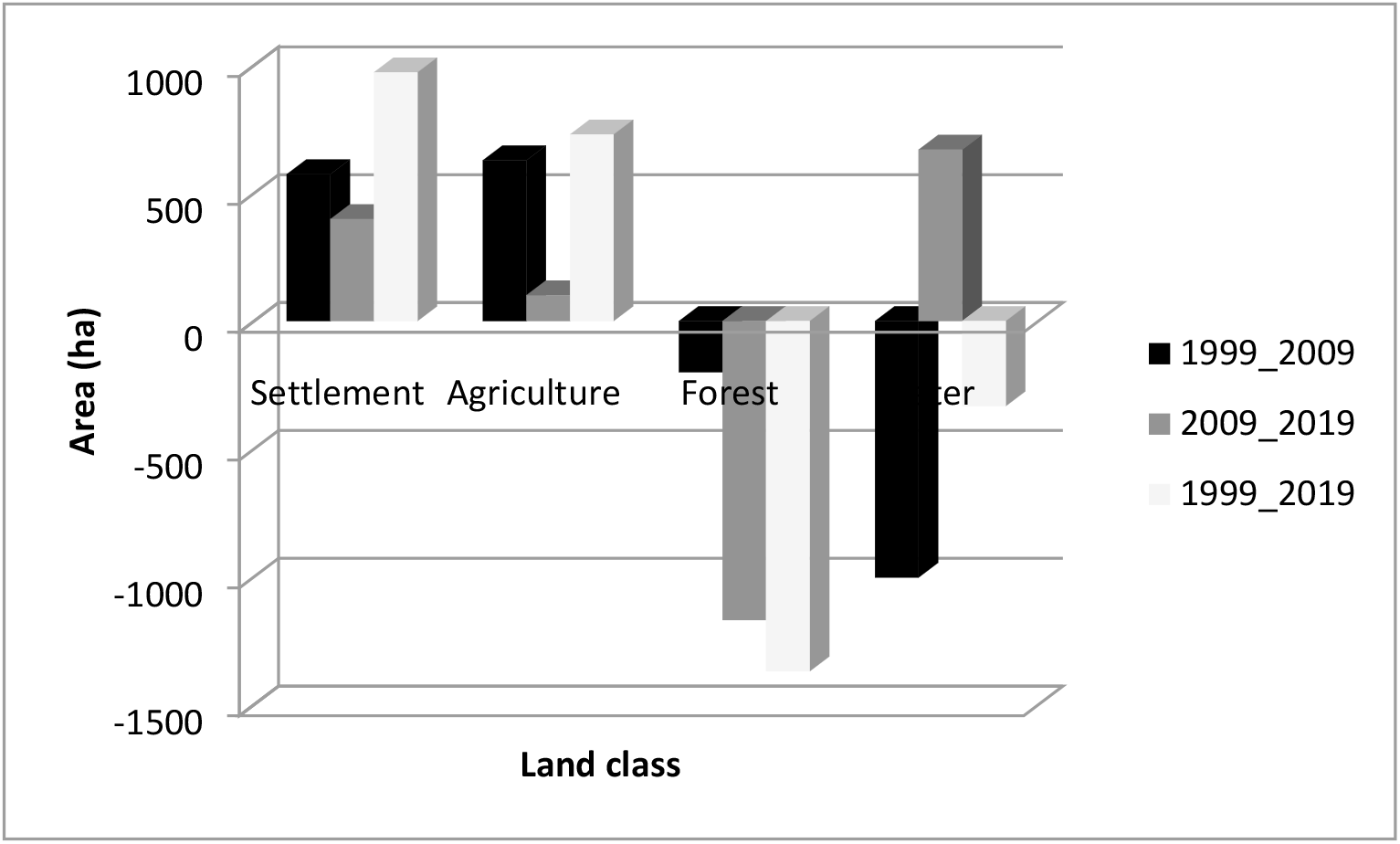
Change detection of the study area

### 3.4 Overall accuracy assessment (1999, 2009 and 2019)

The accuracy of image classification was checked with an accuracy matrix using 140, 173 and 158 randomly selected control points, respectively. The accuracy assessment was performed using land-use maps, ground truth points and Google Earth. Three periods (1999, 2009 and 2019) land use classification have shown, user’s accuracy and producer’s accuracy are greater than 85%, as well the overall accuracy of 92.86%, 94.22% and 94.3% (Table 7,8 and 9), respectively (Table 6, 7 and 8). These values indicate the LAND SAT images and the methodologies used were so accurate. The Kappa coefficient was also calculated, with a value of K= 0.9, which indicates that the classification is almost perfect since it is greater than 0.8. [45] argued that overall accuracy values greater than 0.8 indicate in the Landsat and the methodologies used to have high accuracy.

**Table 6:**
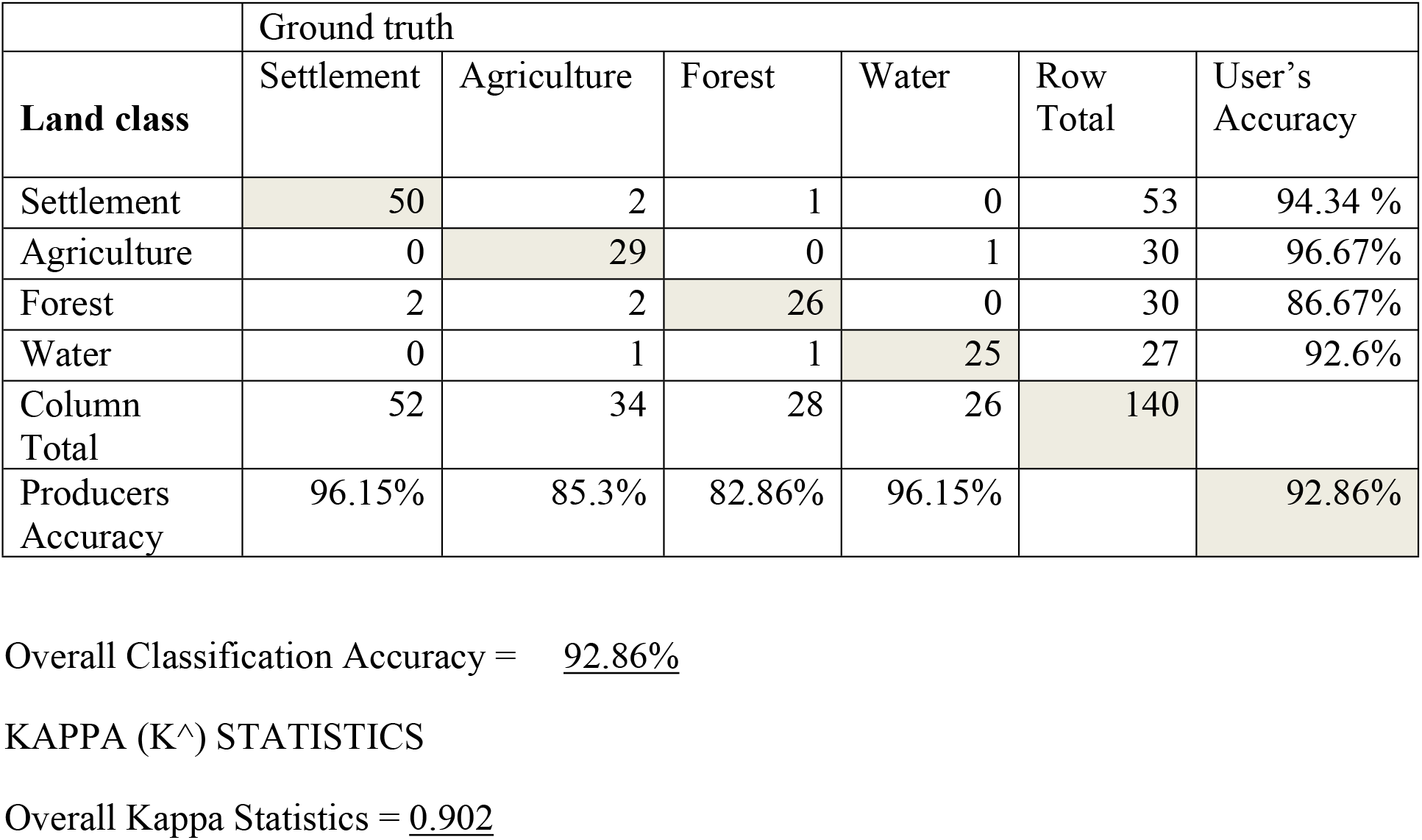
Overall accuracy 0f the study area (1999)

|  | Ground truth |  |  |  |  |  |
| --- | --- | --- | --- | --- | --- | --- |
| **Land class** | Settlement | Agriculture | Forest | Water | Row Total | User's Accuracy |
| Settlement | 50 | 2 | 1 | 0 | 53 | 94.34 % |
| Agriculture | 0 | 29 | 0 | 1 | 30 | 96.67% |
| Forest | 2 | 2 | 26 | 0 | 30 | 86.67% |
| Water | 0 | 1 | 1 | 25 | 27 | 92.6% |
| Column Total | 52 | 34 | 28 | 26 | 140 |  |
| Producers Accuracy | 96.15% | 85.3% | 82.86% | 96.15% |  | 92.86% |

Overall Classification Accuracy = 92.86%
KAPPA (K<sup>^</sup>) STATISTICS
Overall Kappa Statistics = 0.902

**Table 7:** Overall accuracy 0f the study area (2009)

|  | Ground truth |  |  |  |  |  |
| --- | --- | --- | --- | --- | --- | --- |
| **Land class** | Settlement | Agriculture | Forest | Water | Row Total | User's accuracy |
| Settlement | 47 | 2 | 1 | 0 | 50 | 94% |
| Agriculture | 1 | 60 | 2 | 0 | 63 | 95.25% |
| Forest | 1 | 1 | 36 | 0 | 38 | 94.74% |
| Water | 0 | 1 | 1 | 20 | 22 | 90.91% |
| Column Total | 49 | 64 | 40 | 20 | 173 |  |
| Producers Accuracy | 95.92% | 93.75% | 90% | 100% |  | 94.22% |

Overall Classification Accuracy = 94.22%
KAPPA (K<sup>^</sup>) STATISTICS
Overall Kappa Statistics = 0.92

**Table 8:** Overall accuracy 0f the study area (2019)

|  | Ground truth |  |  |  |  |  |
| --- | --- | --- | --- | --- | --- | --- |
| **Land class** | Settlement | Agriculture | Forest | Water | Row Total | User's accuracy |
| Settlement | 54 | 2 | 1 | 0 | 57 | 94.74% |
| Agriculture | 1 | 40 | 1 | 0 | 42 | 95.25% |
| Forest | 1 | 1 | 30 | 0 | 32 | 93.75% |
| Water | 0 | 1 | 1 | 25 | 27 | 92.59% |
| Column<br>Total | 56 | 44 | 33 | 25 | 158 |  |
| Producers<br>Accuracy | 96.43% | 90.91% | 90.91% | 100% |  | 94.3% |

Overall Classification Accuracy = **94.3%**
KAPPA (K<sup>^</sup>) STATISTICS
Overall Kappa Statistics = **0.922**

**Table 9:** NDVI result of the study area

| Statistics | 1999 | 2009 | 2019 |
| --- | --- | --- | --- |
| $NDVI = \frac{(NIR - R)}{NIR + R}$ | | | |
| **Low** | -0.42 | -0.28 | -0.37 |
| **High** | 0.73 | 0.64 | 0.51 |
| **Mean** | 0.18 | 0.059 | -0.21 |
| **SD** | 0.11 | 0.095 | 0.11 |

### 3.5 Normalized difference vegetation index (NDVI)

The statistics and visual observation of the NDVI images over three successive periods (1999, 2009 and 2019) showed that major land cover changes have taken in the study area (Figure 7). The threshold value of NDVI was approximately 0.73 (Figure 6). The pixels having an NDVI value above the threshold were identified as vegetated areas, while low NDVI values represented non-vegetated areas. For non-vegetated areas, we found that the water bodies were represented by low NDVI values, ranging from -0.28 to -0.42, while the pixels having NDVI values in the range of 0.51 to 0.73 were considered as vegetation cover areas (Table 9). NDVI analysis has proven that there had been changes in vegetation cover between 1999 and 2019 images and higher values were recorded in the period 1999 in the study area.

**Figure 6:**
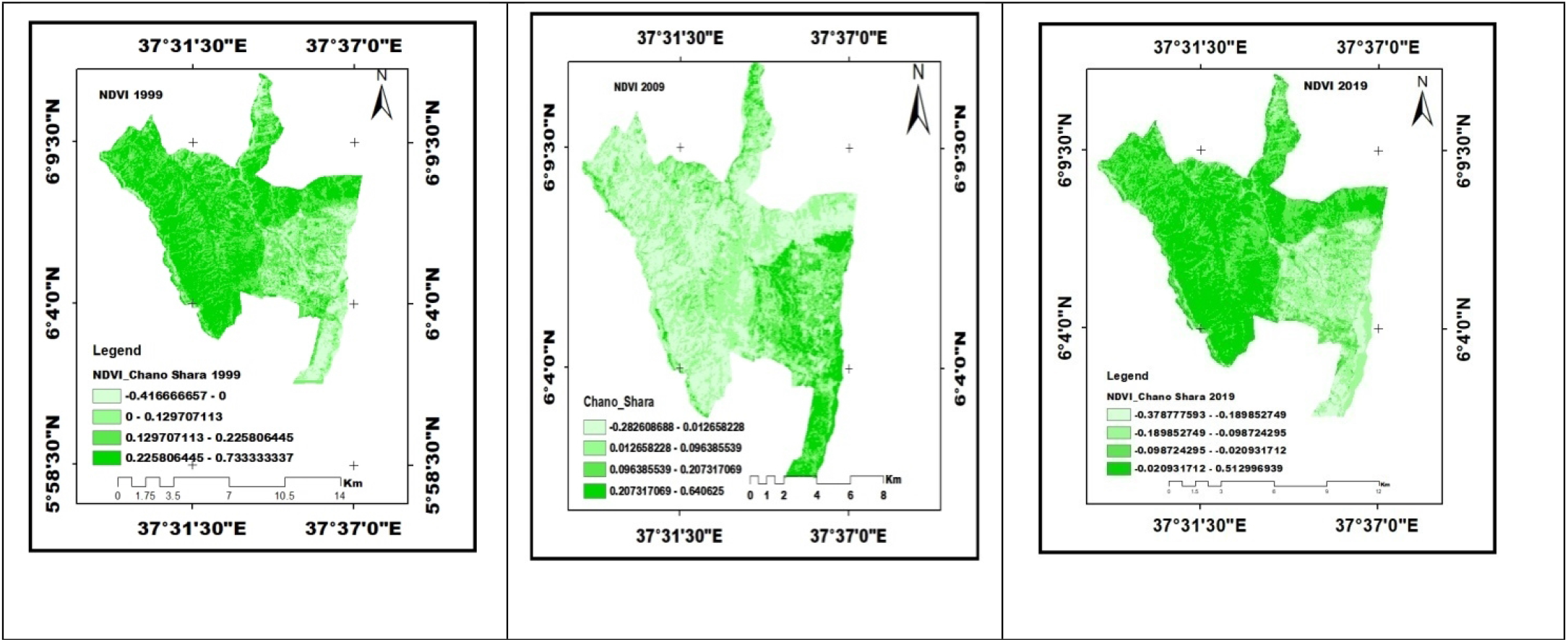
NDVI of the study area from 1999_2019

### 3.6 Drivers of LULC changes

The results of FGD and KII reveal the five major direct driving forces (Table 10). Among these, agricultural expansion account, expansion of settlement and Fuelwood collection and Charcoal production take large shares.

**Table 10:** Proximate causes of LULC changes

| No | Driver | Frequency | % | Rank |
| --- | --- | --- | --- | --- |
| 1 | Fuelwood collection, tree cutting and Charcoal production | 8 | 22.9 | 3 |
| 2 | Agricultural expansion | 11 | 31.4 | 1 |
| 3 | Expansion of settlement | 9 | 25.7 | 2 |
| 4 | Fire | 2 | 5.7 | 5 |
| 5 | Overgrazing | 5 | 14.3 | 4 |
|  | Total | 35 | 100 |  |

The demographic data of the study area over the past three decades has revealed that population pressure ranked as the top cause of LULC changes (Table 11) [46]. The work of Lambin *et al* (2003) show that impact human population pressure is causing the accelerated conversion of natural habitats into agricultural and settlement areas to meet the mounting demand for food and housing. In Ethiopia, resettlement and villagization programs during the Military Government had made a significant contribution to the expansion of settlements and agriculture. Due to the low policy enforcing capacity of the then government landless farmers cleared forests and occupied as much land as possible to increase the chances of land ownership.

**Table 11:**
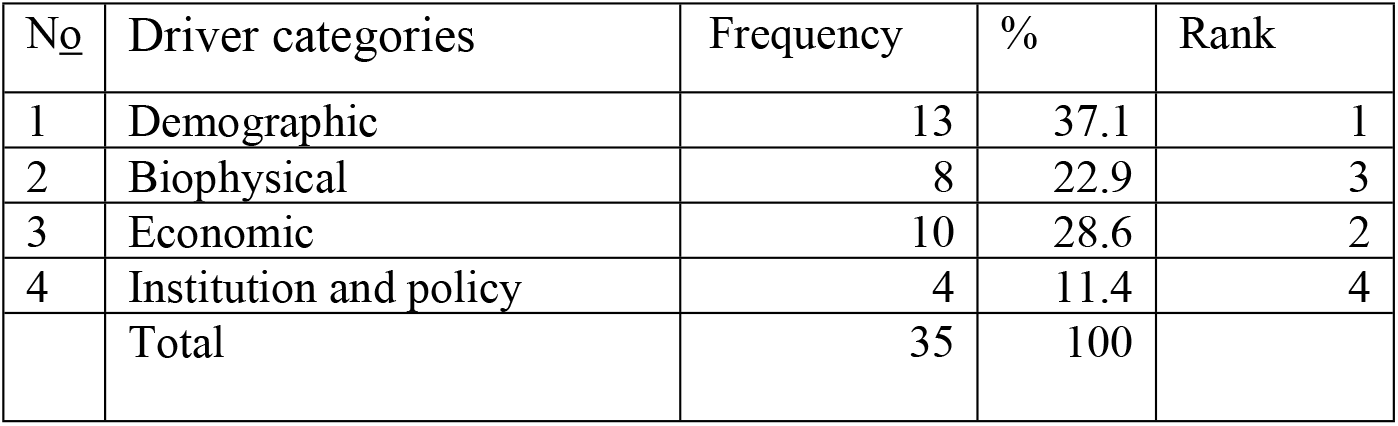
Underlying causes of LULC changes

### 3.7 CONCLUSION and RECOMMENDATIONS

There were four land classes in the study area including forest, farmland, settlement and water bodies and wetlands. The changes observed in 2009 and 2019 were more rapid than that in 1999 the expansion of small-scale irrigated farmlands for fruit and vegetable production. Field observations, KII and focus group discussant confirmed that the main cause of LULC changes in the study area was the expansion of farmland and settlement. On the other hand, demographic, economic and biophysical conditions were indirect driving forces of LULC changes.

Linking participatory forest management with an institution and strong monitoring policies, green legacy and creating awareness to local people is hopped to improve the current status forest biodiversity and environment of the study area. Furthermore, the land use policy and environmental rehabilitation policies of the country need to be revised to include biodiversity hotspots and sequestration of carbon for carbon trading. The environmental trade-offs of fruit and vegetable productions that fetch good economic income must be mitigated through payment for ecosystem services that can be channeled for payment to the workforce involved in green legacy and environmental rehabilitation. Furthermore, promoting none agricultural economy to the jobless youths and creating forest reserved areas with a buffer zone of the study area.

## ACKNOWLEDGEMENT

First of all, I would like to express my special thanks to almighty God who helped me in all my success. I am also very grateful to my supervisors Sebsebe Demissew, Zerihun Woldu and Serekebirhan Takele for their consistent and stimulating advice, valuable suggestions, constructive criticism, reading of the manuscript without his sincere collaborations the work would not have been completed. Moreover, I thank Arba Minch University for provided financial help throughout my research work and next, my gratitude goes to Biodiversity Conservation and Research Centre, School of Graduated studies and Biology Department for all support.

